# DISPOT: A simple knowledge-based protein domain interaction statistical potential

**DOI:** 10.1101/525535

**Authors:** Oleksandr Narykov, Dmitry Korkin

**Affiliations:** Department of Computer Science, Worcester Polytechnic Institute, Worcester, MA, USA; Bioinformatics and Computational Biology Program, Worcester Polytechnic Institute, Worcester, MA, USA

## Abstract

**Motivation:** The complexity of protein-protein interactions (PPIs) is further compounded by the fact that an average protein consists of two or more domains, structurally and evolutionary independent subunits. Experimental studies have demonstrated that an interaction between a pair of proteins is not carried out by all domains constituting each protein, but rather by a select subset. However, finding which domains from each protein mediate the corresponding PPI is a challenging task.

**Results:** Here, we present <u>D</u>omain <u>I</u>nteraction <u>S</u>tatistical <u>POT</u>ential (DISPOT), a simple knowledge-based statistical potential that estimates the propensity of an interaction between a pair of protein domains, given their SCOP family annotations. The statistical potential is derived based on the analysis of more than 352,000 structurally resolved protein-protein interactions obtained from DOMMINO, a comprehensive database on structurally resolved macromolecular interactions

**Availability and implementation:** DISPOT is implemented in Python 2.7 and packaged as an open-source tool. DISPOT is implemented in two modes, basic and auto-extraction. The source code for both modes is available on Github: (https://github.com/KorkinLab/DISPOT) and standalone docker images on DockerHub: (https://cloud.docker.com/u/korkinlab/repository/docker/korkinlab/dispot).

## Introduction

Large-scale characterization of protein-protein interactions (PPIs) using high-throughput interactomics approaches, such as yeast-two-hybrid (Y2H) and tandem-affinity purification/mass spectrometry (TAP/MS) approaches (Gavin, et al., 2002; Rolland, et al., 2014), have provided the scientists with the new insights of the cell functioning at the systems level and allowed to better understand the molecular machinery underlying complex genetic disorders (Barabasi and Oltvai, 2004; Cui, et al., 2015; Mitra, et al., 2013). Structural studies of protein-protein interactions have revealed that a protein-protein interaction is often carried out by smaller structural protein subunits, protein domains (Ekman, et al., 2005; Jin, et al., 2009; Vogel, et al., 2004). Roughly two-thirds of eukaryotic and more than one-third of prokaryotic proteins are estimated to be multi-domain proteins (Ekman, et al., 2005), and thus it is not surprising that ~ 46% of structurally resolved interactions are domain-domain interactions (Kuang, et al., 2016). A high-throughput breakdown of the interactome at this, domain-level, resolution is a much more experimentally challenging task, currently unfeasible at the whole-system level and requiring computational methods to step in (Deng, et al., 2002; Finn, et al., 2004; Ohue, et al., 2014; Segura, et al., 2015).

Here, we present a simple knowledge-based <u>D</u>omain <u>I</u>nteraction <u>S</u>tatistical <u>POT</u>ential (DISPOT), a tool that leverages the statistical information on interactions shared between the homologous domains from structurally defined domain families. The knowledge-based potentials are extracted form our comprehensive database on structurally resolved macromolecular interactions, DOMMINO (Kuang, et al., 2016). Our statistical potential can be integrated into protein-protein interaction prediction methods that deal with multidomain proteins by ranking all possible pairwise combinations of domain interactions between the two or more proteins.

## Methodology

The development of DISPOT is driven by several observations. First, an average interaction between a pair of proteins is not carried out by all domains constituting each protein, but only by a select subset. Indeed, each domain has its unique structure and biological function and may not be designed to interact with a particular domain from another protein (Banappagari, et al., 2010; Shimizu, et al., 2016). Second, the domain-domain interactions often share homology: when two homologous domains interact with their partners, these partners frequently also share the homology with each other (Kuang, et al., 2016). Thus, one can introduce the domain-domain interaction propensity in terms of the frequency of domain-domain interactions between the two domain families. Lastly, the propensity of domain-domain interaction is expected to vary across different families, thus allowing to provide the finer resolution of the protein-protein interaction network.

The quantification of the odds for a domain from one domain family to interact with a domain from another family is defined in this work as a knowledge-based statistical potential. Statistical potentials are widely used in biophysical applications, often for characterizing the residue contacts between the protein chains (Krüger, et al., 2014; Lu, et al., 2003). One of the main applications of the residue-level statistical potentials is in protein docking (Kozakov, et al., 2006). Our domain-domain statistical potential complements the residue-level potentials by considering structural units from the higher-level of protein structure hierarchy and requiring no structural information about the protein domains. Specifically, the input for DISPOT includes the protein sequences of the two proteins interacting with each other.

First, the domain architecture of each protein is obtained. To do so, a region of the protein sequence is annotated to a family of homologous domains. For the definition of domain families, we leverage the Structural Classification of Proteins (SCOP) family-level classification (Andreeva, et al., 2004). SCOP represents a structure-based hierarchical classification of relationships between protein domains or single-domain proteins with ‘family’ being the first level of SCOP classification and ‘superfamily’ being the second level. Protein domains from the same SCOP family are evolutionary closely related and often share the same function. Since a protein with no structural information cannot be directly annotated by SCOP, we use SUPERFAMILY (Gough and Chothia, 2002), a Hidden Markov Model (HMM) based approach that maps regions of a protein sequence to one or several SCOP families or superfamilies. SUPERFAMILY allows us to cover a substantial subset of known proteins: the HMM coverage at the protein sequence and amino acid levels for the UniProt database were reported at 64.73% and 58.78% respectively in 2014 (Oates, et al., 2014).

Second, for each pair of SCOP families we count a number of non-redundant protein-protein interactions between the members of these families that have been experimentally determined. Our source of data is DOMMINO (Kuang, et al., 2016; Kuang, et al., 2011) a comprehensive database of structurally resolved macromolecular interactions. It contains information about interactions between the protein domains, interdomain linkers, terminal sequences, and protein peptides. In this work, we use exclusively domain-domain interactions because the data about this type of interactions is the most abundant. To remove redundancy in the data, we use ASTRAL compendium (Brenner, et al., 2000) which is integrated into the SCOPe database (Fox, et al., 2013). From ASTRAL, we obtain a set of domains, where each domain shares less than 95% sequence identity to any other domain in the set. This set is then used to determine pairs of redundant domain-domain interactions in the original DOMMINO dataset. Two domain-domain interactions are determined as redundant if both corresponding pairs of domains share 95% or more sequence identity. For each pair of redundant domain-domain interactions, one interaction is randomly removed. The process continues until no pair of redundant interactions can be detected.

Third, for each domain family from each protein, a statistical potential is calculated (Fig. 1A). There are two types of statistical potentials introduced in this work: (1) calculated for a domain from a specific domain family, and (2) calculated for a pair of domains, one domain from each of the two interacting proteins. The statistical potential ***P_i_*** for a single domain ***D_i_*** is calculated based on the total number of interactions ***N_Di_*** extracted from the non-redundant DOMMINO dataset for the specific SCOP family this domain belongs to. The statistical potential ***P_ij_*** for a pair of domains, ***D_i_*** and ***D_i_***, is calculated based on the total number of occurrences ***N_ij_*** of the interactions between all domains from the same two SCOP families as ***D_i_*** and ***D_ij_***. Those numbers are then transformed into probabilities as follows:

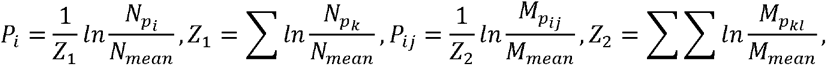

where *N_mean_* is an average number of interactions for a domain family and *M_mean_* is an average number of interactions for a pair of domain families, both calculated from the non-redundant DOMMINO set.

**Figure 1.**
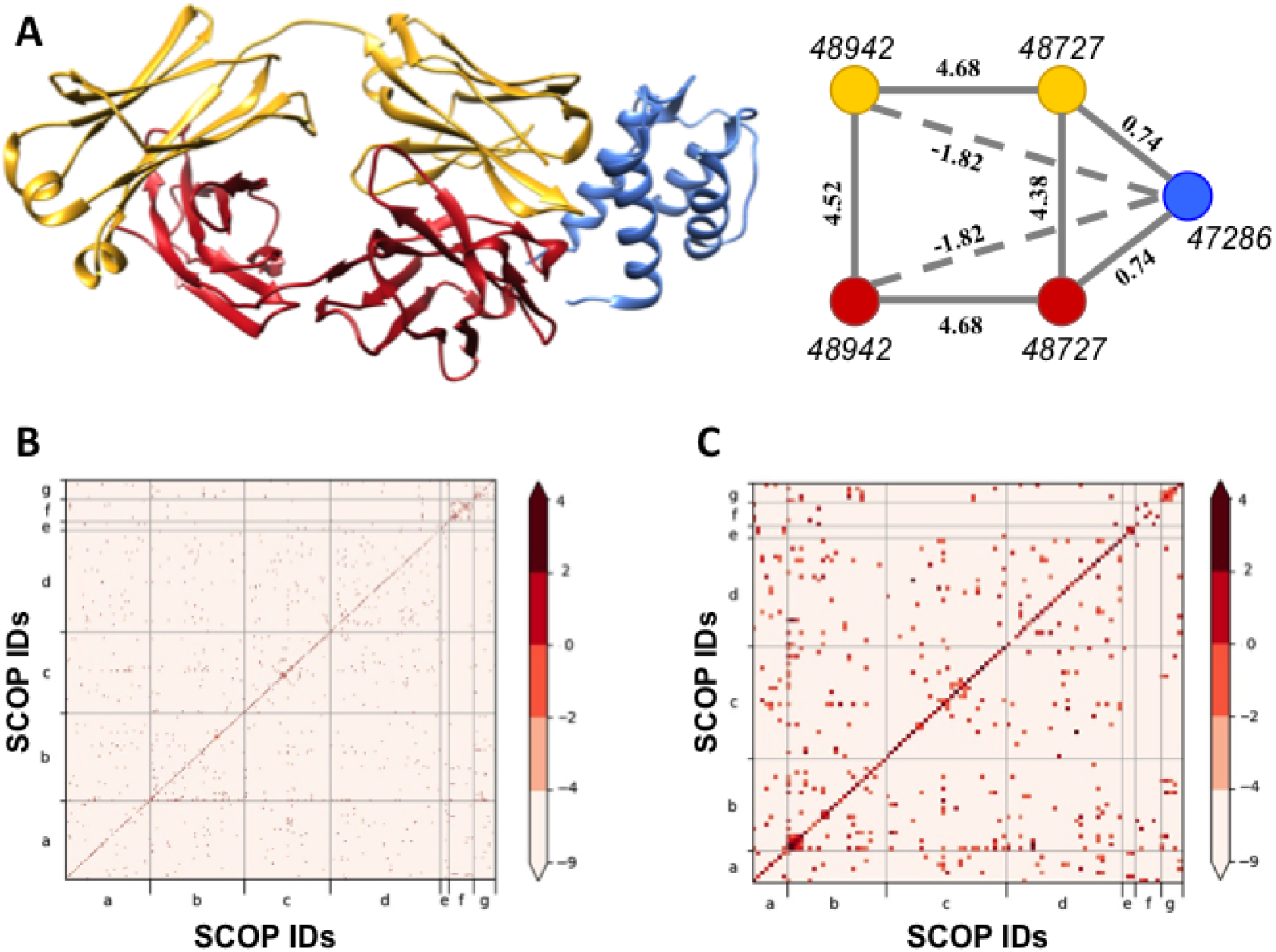
DISPOT statistical potential and its application. **A**. A crystal structure (left) of the protein complex between CNTO607 Fab human monoclonal antibody (yellow and red colors denote two different chains) and interleukin-13 (IL-13), and the corresponding domain-domain interaction network (right). Shown in italics are SCOP Family IDs, and in bold are DISPOT values for the corresponding interactions. Nodes colored with the same color belong to the same chain. Solid lines connecting nodes correspond to the physical interaction, while dashed lines connect nodes corresponding to the protein domains that do not physically interact. **B**. A heatmap showing DISPOT values calculated for each pair of SCOP families, where only potentials for pairs of SCOP families with 5 and more non-redundant interactions are plotted. The families are grouped based on the SCOP class (a-g) and are ordered within each fold based on their IDs. **C**. A heatmap showing DISPOT values calculated for each pair of SCOP families, where the pixel size for each entry was increased compared to that one in **B**.

Overall, we have analyzed and summarized interactions from 3,619 SCOP family pairs that were extracted from 352,199 PPIs. In total, domains from 1,384 SCOP families were characterized that form domain-domain interactions in 1,384 ‘homo-SCOP’ interaction pairs (*i.e*., both domains are annotated with the same SCOP family) and 2,235 ‘hetero-SCOP’ pairs (Fig. 1B,C). The analysis of the calculated statistical potentials showed a wide diversity across different families.

## Implementation and usage

The basic mode is implemented in Python with the dependency on packages *pandas* and *numpy*. It takes SCOP identifiers (IDs) for either ‘family’ (fa) or ‘superfamily’ (sf) hierarchy levels as an input and produces statistical potential for corresponding pair of domains. Switching between the SCOP levels is implemented in command line option *sf*. One of the possible input options is a command line option *domains*, which provides a list of space-separated SCOP identifiers. Based on this list, the program produces all possible unique pairwise combinations of identifiers and the corresponding statistical potentials. Option *max* produces the highest value of statistical potential for a selected domain and a SCOP ID for the corresponding interaction domain partner. Option *output* specifies the output file. If no file path is specified, then program opens a console output prompting a user to input the data. A detailed description of all acceptable input formats and options is available in README file and help menu of the main script *dispot.py*.

The auto-extraction version relies on the SUPERFAMILY models and scripts and HMMER program for extracting the corresponding SCOP IDs for either family or superfamily levels of hierarchy. The Perl programming language interpreter is an additional dependency. HMMER is compatible with the major linux distributions (it has been tested on Ubuntu 16.04 and Alpine 3.7). Windows users are advised to use the docker image. The main script is *dispot.py*, and it includes several options: *fasta_folder*– to specify a path to the folder with FASTA files; *output_folder –* to specify a path to the results; and *max*– to substitute the regular output of all pairwise statistical potentials with the highest statistical potential for a given domain family and a SCOP ID of the interaction partner on which this value is achieved. Additional script *batch_process.py* provides almost the same functionality, except it uses the default locations: *./data* for the input and *./data/results* for the output. For each FASTA sequence, we extract a SUPERFAMILY-derived SCOP ID and the location(s) of the corresponding domain on the protein sequence. It is stored in the *./tmp* folder and is available until the next run of any of the scripts mentioned in this section. The data are stored in the Python dictionary objects serialized by package *pickle*.

